# Cotyledon Peeling Method for Passion Fruit Protoplasts: a versatile cell system for transient gene expression in passion fruit (*Passiflora edulis*)

**DOI:** 10.1101/2023.06.08.542798

**Authors:** Linxi Wang, Haobin Liu, Peilan Liu, Guanwei Wu, Wentao Shen, Hongguang Cui, Zhaoji Dai

## Abstract

Passion fruit (*Passiflora edulis*) is a perennial evergreen vine that grows mainly in tropical and subtropical regions due to its nutritional, medicinal and ornamental values. However, the molecular biology study of passion fruit is extremely hindered by the lack of an easy and efficient method for transformation. The protoplast transformation system plays a vital role in plant regeneration, gene function analysis and genome editing. Here, we present a new method (‘Cotyledon Peeling Method’) for simple and efficient passion fruit protoplast isolation using cotyledon as the source tissue. A high yield (2.3 × 10^7^ protoplasts per gram of fresh tissues) and viability (76%) of protoplasts were obtained upon incubation in the enzyme solution [1% (w/v) cellulase R10, 0.25% (w/v) macerozyme R10, 0.4 M mannitol, 10 mM CaCl_2_, 20 mM KCl, 20 mM MES and 0.1% (w/v) BSA, pH 5.7] for 2 hours. In addition, we achieved high transfection efficiency of 83% via the polyethylene glycol (PEG)-mediated transformation with a green fluorescent protein (GFP)-tagged plasmid upon optimization. The crucial factors affecting transformation efficiency were optimized as follows: 3 μg of plasmid DNA, 5 min transfection time, PEG concentration at 40% and protoplast density of 100 × 10^4^ cells/ml. Furthermore, the established protoplast system was successfully applied for subcellular localization analysis of multiple fluorescent organelle markers and protein-protein interaction study. Taken together, we report a simple and efficient passion fruit protoplast isolation and transformation system, and demonstrate its usage in transient gene expression for the first time in passion fruit. The protoplast system would provide essential support for various passion fruit biology studies, including genome editing, gene function analysis and whole plant regeneration.

## 1 Introduction

Passion fruit (*Passiflora edulis*), also known as passiflora, is a perennial evergreen vine that grows mainly in tropical and subtropical regions due to its nutritional, medicinal and ornamental values. It belongs to the genus *Passiflora*, the largest genus in the Passifloraceae family, and originated in South America (Fischer and Rezende, 2008; Xia et al., 2021; Yu et al., 2021; Phong et al., 2022). Passion fruit has been gaining popularity as a flavor in both drinks and foods as it’s rich in vitamins, antioxidants, and plant compounds that could benefit human health. In China, it is widely distributed in several provinces with warm and humid climates, including Hainan, Taiwan, Fujian, Yunnan, Guangxi and Guizhou. In the past decades, passion fruit plantations soared in China as an economic investment (Yu et al., 2021). However, the molecular biology study of passion fruit falls far behind. It was not until 2021 that a chromosome-scale genome assembly of passion fruit (*Passiflora edulis* Sims) was just reported (Xia et al., 2021). This work enables several documentations for the identification of passion fruit genes, including *eceriferm* (*CER*), *aquaporin* (*AQP*), *β-Ketoacyl-CoA synthase* (*KCS*), *peroxidase* (*POD*), *lateral organ boundary domain* (*LBD*), and *lipoxygenase* (*LOX*) gene families (Rizwan et al., 2022a; Song et al., 2022; Rizwan et al., 2022b; Liang D. et al., 2022; Liang J. et al., 2022; Huang et al., 2022). However, one of the major barriers in gene functional studies is the lack of a simple and efficient transformation system in passion fruit.

Plant transformation is a fundamental and powerful tool for molecular plant biology studies. There are two types of plant transformation: stable transformation and transient transformation. The *Agrobacterium*-mediated transformation has been extensively used to generate transgenic plants. For passion fruit, G. Manders and his colleagues generated the first transgenic passion fruit plants by *Agrobacterium tumefaciens*-mediated transformation using leaf as explant in 1994 (Manders et al., 1994). After that, many groups worldwide have reported the successful in vitro regeneration and *Agrobacterium*-mediated transformation of passion fruit (Trevisan et al., 2006; Correa et al., 2015; Rizwan et al., 2021) The *Agrobacterium*-mediated transformation method represents a powerful platform for transgenic passion fruit lines, but exhibits the disadvantages of time-consuming, relatively low transformation efficiency and inconvenience for large-scale gene functional analyses. Alternatively, transient gene expression represents a rapid and high-throughput method for gene functional studies (Zhao et al., 2016). The most common transformation methods for transient gene expression include protoplast transformation, biolistic bombardment and *Agrobacterium tumefaciens*-mediated transformation (Pitzschke and Persak, 2012). Although protoplast transformation has been widely used for transient gene expression analysis in a large number of plants, including Arabidopsis, maize and rice, there is currently no report for transient gene expression in passion fruit.

Plant protoplasts are the plant single cells excluding the rigid cell wall, and serve as a versatile single-cell-based system for transient gene expression (Davey et al., 2005; Dai and Wang, 2022; Wang et al., 2023). Foreign plasmid DNA of interest can be easily delivered into protoplasts typically by polyethylene glycol (PEG)-mediated transformation, electroporation and microinjection (Yoo et al., 2007; Priyadarshani et al., 2018; Dai and Wang, 2022). Since the first report of successful plant protoplasts isolation from tomato seedlings in 1960 by Cocking (Cocking, 1960), the protoplast transformation system has been extensively applied in transient gene expression such as intracellular localization, protein-protein interaction and genome editing studies (Davey et al., 2005; Yu et al., 2017; Priyadarshani et al., 2018). This system relies on efficient protoplast isolation from plant tissue and successful downstream transformation. Multiple parameters can influence the transformation efficiency of PEG-mediated transformation, including incubation time, PEG concentration, protoplast density and plasmid amount.

The aim of this study was to establish a platform for passion fruit transient gene expression. Owing to the wealth of knowledge of plant protoplast for transient gene expression, the protoplast system would likely provide a feasible way for transformation in passion fruit. Toward this goal, we report here a simple method of using peeled seedling cotyledon for passion fruit protoplast isolation and achieved high transfection efficiency of 83% upon optimization. The simple and efficient protoplast transformation system can be broadly applied in transient gene expression, illustrated by intracellular localization analysis of multiple fluorescent markers as well as protein-protein interaction study. This is the first study to describe protoplast transformation system for transient gene expression in passion fruit. The significance of these results for promoting the molecular biology study of passion fruit is discussed.

## 2 Materials and Methods

### 2.1 Plant material and growth conditions

Passion fruit (*Passiflora edulis*) seeds were placed in the soil at a depth of 1-2 cm and germinated after 7-10 days with 16 hours light/8 hours dark regime at 28±2 °C. The cotyledons were fully expended after approximately 7 days upon germination and could be readily used for protoplast isolation.

### 2.2 Plasmid construction and preparation

The green fluorescent protein (GFP)-expressing plasmid pGreen0029-GFP was a gift from Dr. Ji Li of Nanjing Agriculture University, China. H2B-RFP and ER-mCherry-HDEL were obtained from Dr. Guanwei Wu of Ningbo University, China.

YN-CP and YC-CP are constructed by the following methods: The full length of *CP* gene (GenBank: MG944249.2) was amplified from the infectious cDNA clone of telosma mosaic virus (TelMV) using PCR with gene-specific primers (CP-F: 5’ GGGGACAAGTTTGTACAAAAAAGCAGGCTTCATGTCTGGAAAGGTTGATGATG3’; CP-R: 5’GGGGACCACTTTGTACAAGAAAGCTGGGTCCTGCACAGAACCTACTCC3’) and subcloned into the Gateway entry vector pDONR 221 (Invitrogen) by BP reaction and finally recombined into the destination vector pEarleyGate 201-nYFP and pEarleyGate 202-cYFP vector (Dai et al., 2020) by LR reaction, respectively.

All the constructed plasmids are validated by double digestion and sequencing. Plasmids were extracted using the Maxi Plasmid Kit Endotoxin Free (Geneaid) according to the manufacturer’s instructions.

### 2.3 Passion fruit protoplast isolation

The fully expended cotyledons were attached to the 3M Masking tape with the upper side facing down and peeled away the lower epidermal surface by directly pulling the cotyledons. The peeled cotyledons were further immersed in the enzyme solution [1% (w/v) cellulase R10 (Yakult Pharmaceutical Ind. Co., Ltd., Japan), 0.25% (w/v) macerozyme R10 (Yakult), 0.4 M mannitol, 10 mM CaCl_2_, 20 mM KCl, 20 mM MES and 0.1% (w/v) BSA, pH 5.7] in a Petri dish. The peeled cotyledons were incubated for 2-4 hours with gently shake (40 rpm on a platform shaker) in the dark at room temperature (25 °C) for releasing protoplasts and were followed by raising using the iced W5 (154 mM NaCl, 125 mM CaCl_2_, 5 mM KCl, 5 mM Glucose, 2 mM MES, pH 5.7).

Passion fruit protoplasts were then transferred to round-bottom centrifuge tubes by passing through a 100 μm nylon mesh filter and pelleted for 3 min at 100 g. The pelleted protoplasts were washed with ice-cold W5 once. Re-suspended the protoplasts in 300 μl W5 and rest on ice for 30 min. Finally, resuspend the protoplasts in MMg solution (0.4 M mannitol, 15 mM MgCl_2_, 4 mM MES, pH 5.7) at a density of 100 × 10^4^ cells /ml at room temperature.

### 2.4 Passion fruit protoplast transformation

We adapted the classic PEG-mediated protoplast transformation method for passion fruit protoplast transformation (Dai and Wang, 2022). Specifically, 30 μl protoplasts were added to 3 μg plasmid in a 2 ml round-bottom centrifuge tube and mixed gently. Then, 33 μl freshly made PEG transfection solution (40% (w/v) PEG 4000, 0.1 M CaCl_2_, 0.2 M mannitol) were added. Samples were mixed well by gentle swirling and incubated for a certain time at room temperature. W5 was added to stop the transformation and subjected to centrifugation for 2 min at 100 g. Protoplasts were washed again using W5 and finally stored in 50 μl of W5 at room temperature in the dark.

### 2.5 Protoplast yield calculation and viability assessment

Protoplast yield calculation and viability assessment were performed as described in our previous paper (Wang et al., 2023)

### 2.6 Microscopy

The protoplasts were observed under a fluorescent microscope (OLYMPUS DP80, Japan). For detailed visualization of GFP, organelle markers, BiFC and chloroplast auto-fluorescence, the transformed protoplasts were observed under a confocal microscope (OLYMPUS FV1000, Japan). Excitation wavelengths and emission filters were 488 nm/bandpass 500-520 nm for GFP, 514 nm/band-pass 510-546 nm for YFP, 543 or 563 nm/bandpass 580-620 nm for RFP, and 488 nm/band-pass 680-720 nm for chloroplast auto-fluorescence.

### 2.7 Statistical analysis

Statistical differences between samples were determined by the unpaired two-tailed Student’s t-test and considered significant at P < 0.05. Data are presented as means ± SD of the mean from at least three experiments.

## 3 Results

### 3.1 Isolation of passion fruit protoplast from seedling cotyledons

To establish a simple and efficient protoplast isolation protocol from fresh tissue of passion fruit, the fully expanded cotyledons of a 15-day-old passion fruit seedling were selected for the source material (Fig. 1A, B). Upon peeling off the lower epidermis (Fig. 1C), the cotyledons were incubated with the enzyme solution for 2-4 hours (Fig. 1D, E) and the solution appeared green upon rinsing (Fig. 1F), indicating protoplasts were released. The passion fruit protoplasts were harvested essentially following the protocol developed for Arabidopsis protoplast isolation with a few modifications (Yoo et al., 2007) (Fig. 1G). With the current protocol, a high yield of protoplasts (2.3 × 10^7^ protoplasts per gram of fresh seedling cotyledons) was achieved. The isolated passion fruit protoplast remains round-shape and intact under the light microscope (Fig. 1H). Furthermore, the protoplast viability test revealed that the purified protoplasts remain viable upon fluorescein diacetate (FDA) staining (Fig. 1I). We were able to achieve an average viability of passion fruit protoplast of 76%. These data demonstrated that the seedling cotyledon served as an excellent protoplast source and we have established a simple and efficient protoplast isolation system for passion fruit.

**Figure 1.**
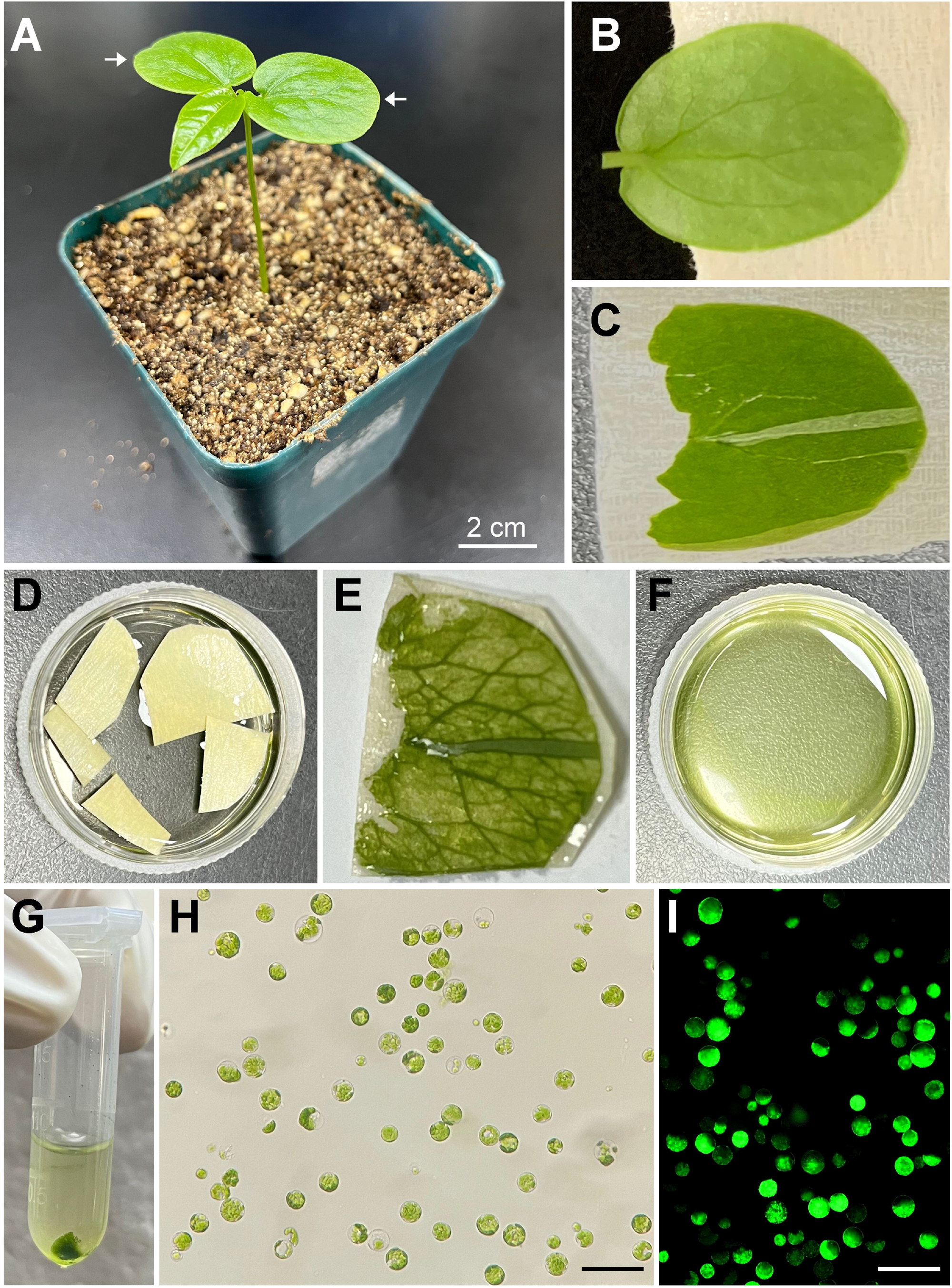
Schematic illustration of passion fruit protoplast isolation from seedling cotyledons. **(A)** A healthy 15-day-old passion fruit seeding plant for cotyledon protoplasts isolation. Arrows indicate the cotyledons to be detached for protoplast isolation. Scale bar, 2 cm; **(B)** Detached cotyledon attached to masking tape with lower side facing up; **(C)** Cotyledon with the lower epidermal surface been pulled away; **(D)** Peeled cotyledon sections incubated with the enzyme solution in a petri dish; **(E)** Cotyledon after digesting with enzyme solution; **(F)** Released protoplasts in the enzyme solution after digestion; **(G)** Protoplasts pellet upon low-speed centrifugation; **(H)** Intact protoplasts in MMg solution under a light microscope after protoplasts purification. Scale bar, 100 µm; **(I)** Fluorescence micrograph of protoplasts emitting green fluorescence upon FDA staining. Scale bar, 100 µm.

### 3.2 Optimization of PEG-mediated transformation of passion fruit protoplasts

Next, we adapted the classic polyethylene-glycol (PEG)-mediated technique (Dai and Wang, 2022) for passion fruit protoplast transformation. A green fluorescent protein (GFP)-tagged plasmid driven by the constitutive CaMV35S promoter (pGreen0029-GFP, 6.0 kb) was used to deliver into passion fruit protoplast to determine the transformation efficiency using a fluorescent microscope. To obtain relatively higher transformation efficiency, various transformation parameters, including plasmid DNA amount, protoplast density, PEG 4000 concentration and incubation time were optimized accordingly.

#### a) Effect of plasmid amount on passion fruit protoplasts transfection

The first variable optimized was the plasmid DNA amount of pGreen0029-GFP. A series of plasmid amounts (0.5, 1, 3, and 6 μg, respectively) was applied to identify the optimal amount of DNA required for PEG-mediated transformation. When using 0.5 μg plasmid for transformation, very few protoplasts exhibited green fluorescence under a fluorescent microscope at 24 hours-post transfection (hpt) (Fig. 2A) and the transfection efficiency was only 12% on average (Fig. 2B). By contrast, the transfection efficiency increased to 30% and 74% as the plasmid amount increases to 1 and 3 μg, respectively (Fig. 2B). When using higher plasmid amount of 6 μg, the transfection efficiency remains high (71%), but no significant difference compared with that of 3 μg (p= 0.45) (Fig. 2B). Thus, we conclude that 3 μg of plasmid is the optimal amount per 30 μl protoplasts transformation reaction.

**Figure 2.**
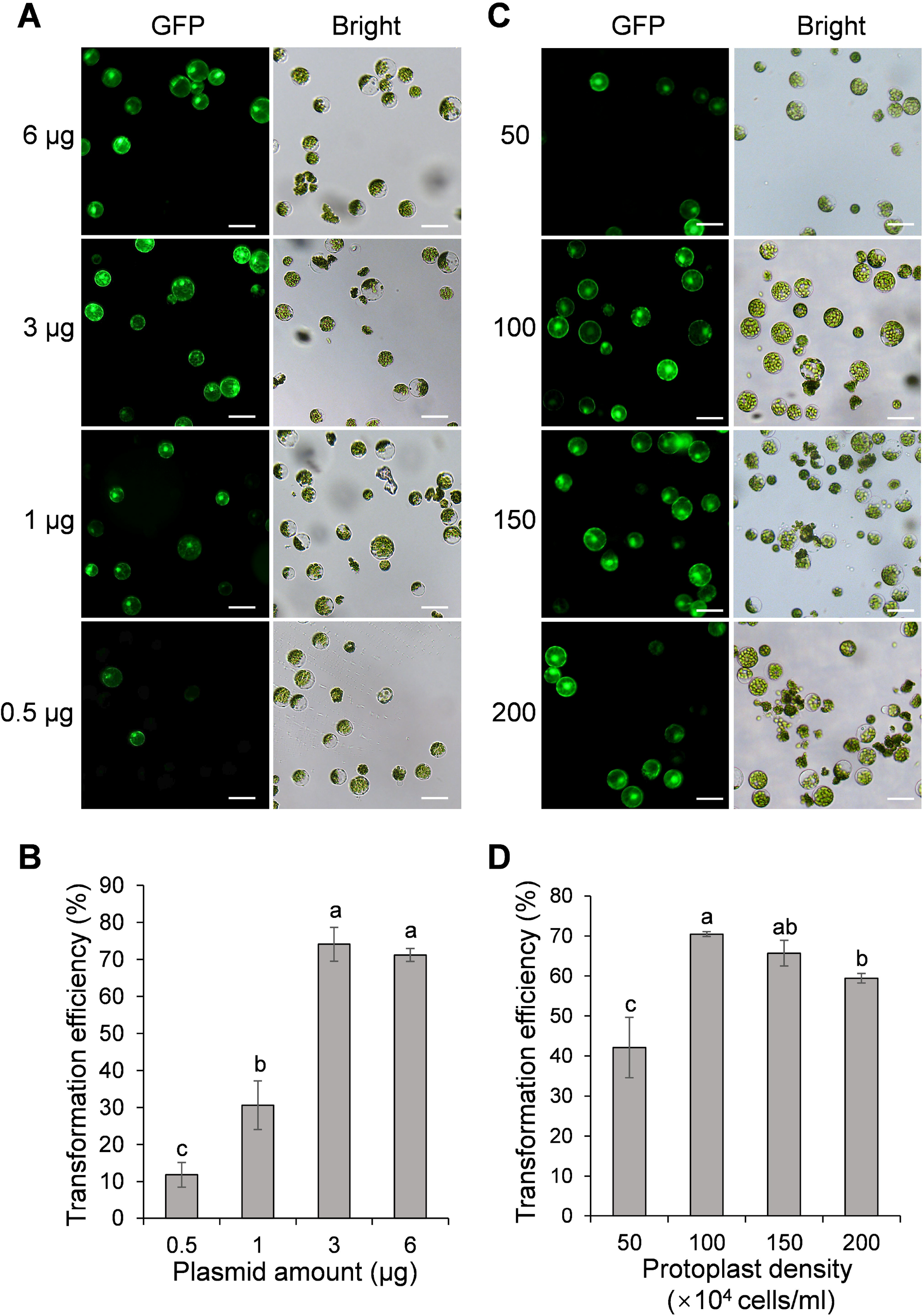
Effect of plasmid amount and PEG 4000 concentration on passion fruit protoplast transfection. **(A)** Microscopic images of passion fruit protoplasts transfected with different amounts of GFP-expressing plasmid (0.5, 1, 3, and 6 μg, respectively). Scale bar, 50 μm. **(B)** The transformation efficiency of passion fruit protoplasts using different amounts of plasmids. Bars represent standard errors. Different letters indicate a statistically significant difference at P < 0.05 among samples according to Duncan’s multiple range tests. **(C)** Microscopic images of passion fruit protoplasts transfected with a GFP-expressing plasmid using varying concentrations of PEG 4000. Scale bar, 50 μm. **(D)** The transformation efficiency of passion fruit protoplasts at various concentrations of PEG4000. Different letters indicate a statistically significant difference at P < 0.05 among samples according to Duncan’s multiple range tests. Bars represent standard errors.

#### b) Effect of protoplast density on transfection efficiency

The second variable optimized was the protoplast density (50, 100, 150 and 200 × 10 ^4^ cells/ml, respectively). We found that low protoplast density at 50 cells/ml produced a transformation efficiency of about only 42% (Fig. 2C, D). By contrast, the highest transformation efficiency (70 ± 1 %) was achieved with the protoplast density at 100 × 10^4^ cells/ml (Fig. 2C, D). Further increasing the protoplast density to 150 × 10^4^ cells/ml did not increase the transformation efficiency (p= 0.06) (Fig. 2D). In addition, a higher protoplast density of 200 × 10^4^ cells/ml resulted in a significant reduction (p= 0.0001) than that of 100 × 10^4^ cells/ml. Hence, we conclude that protoplast density at 100 × 10^4^ cells/ml is optimal in the current protocol.

#### c) Effect of PEG concentration on the transformation of passion fruit protoplasts

We next examined the effect of PEG concentration on the transformation efficiency of passion fruit protoplasts. A series of PEG 4000 concentrations (20, 30, 40, 50 and 60%, respectively) was set to test the protoplast transformation efficiency. Based on the results shown in Fig. 3A, B, transfection efficiency increases significantly as the PEG concentration raise from 20% to 30% and 40%, and reaches a peak of approximately 83% transfection rate at 40% PEG. However, further increasing the PEG concentration from 40% to 50% and 60% gradually reduced the transfection rate. These data demonstrated that the optimal PEG 4000 concentration is 40% for the PEG-based transformation of passion fruit protoplast.

**Figure 3.**
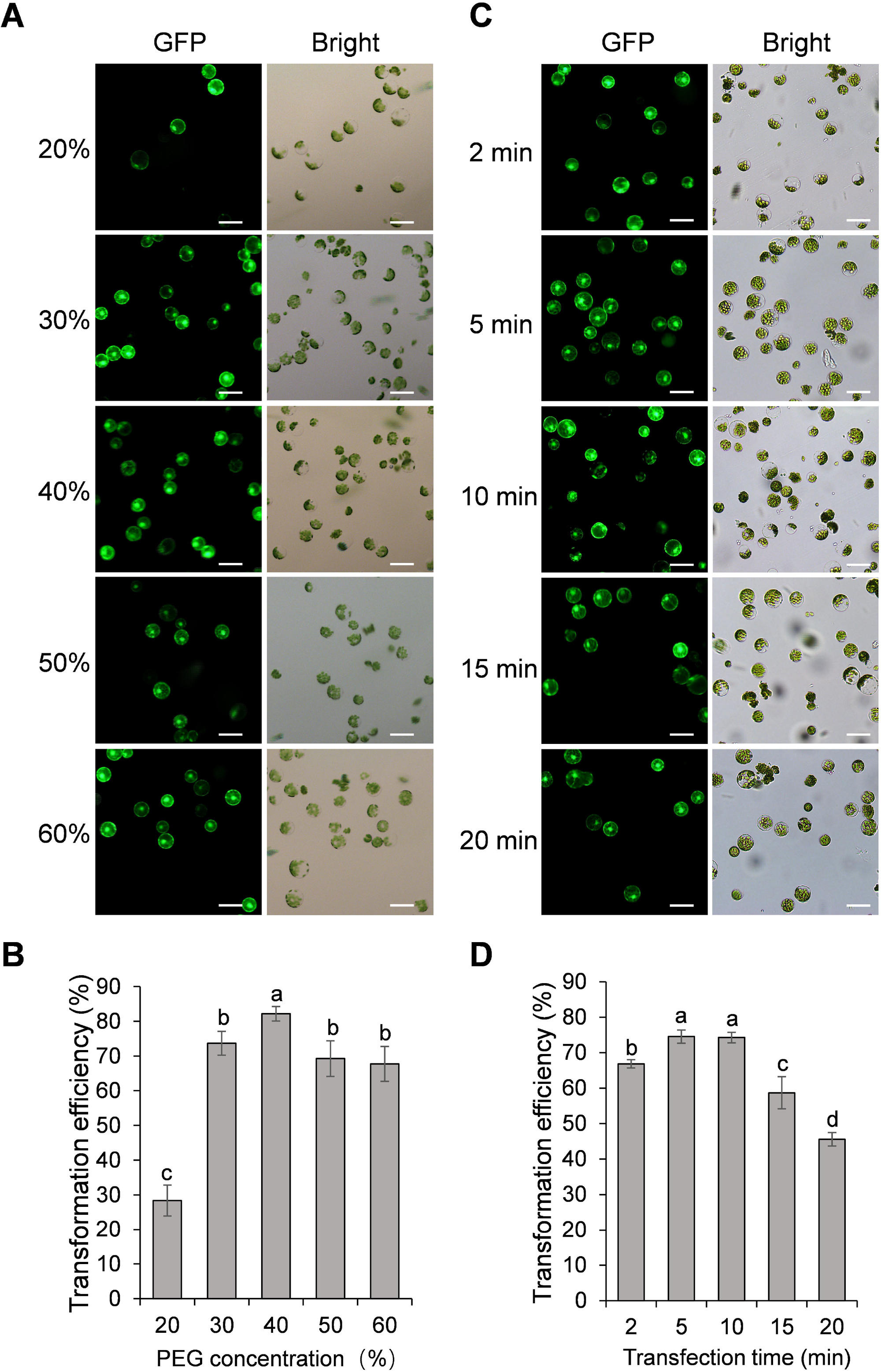
Effect of protoplast density and incubation time on passion fruit protoplast transfection. **(A)** Microscopic images of passion fruit protoplasts transfected with a GFP-expressing plasmid at different densities of protoplast. Scale bar, 50 μm. **(B)** Effect of protoplast density on the transformation efficiency of passion fruit protoplasts. Different letters indicate a statistically significant difference at P < 0.05 among samples according to Duncan’s multiple range tests. Bars represent standard errors. **(C)** Microscopic images of passion fruit protoplasts transfected with a GFP-expressing plasmid at varying incubation time. Scale bar, 50 μm. **(D)** Effect of incubation time on the transformation efficiency of passion fruit protoplasts. Different letters indicate a statistically significant difference at P < 0.05 among samples according to Duncan’s multiple range tests. Bars represent standard errors.

#### d) Effect of incubation time on the transformation efficiency

The last factor to be optimized is the incubation time (2, 5, 10, 15, and 20 min, respectively). As shown in Fig. 3C, D, the highest transfection efficiency (75 ± 2 %) was achieved upon incubation with PEG-calcium transfection solution for 5 min, although there is no significant difference between 5 min and 10 min (P=0.83). Further prolongation of incubation time to 15 min and 20 min significantly decreased the transfection rate to only 59% and 46%, respectively. Therefore, 5 min was taken as the optimal transfection time for PEG-mediated transformation in passion fruit.

Upon all these optimizations, we have successfully established the highly-efficient passion fruit protoplast transformation illustrated by the following points: 1) GFP expression was visualized at 24 hpt under a fluorescence microscope (Fig. 4A). Moreover, confocal laser scanning microscopy (CLSM) revealed that GFP localized both in the nucleus and cytoplasm (Fig. 4B), suggesting GFP was successfully expressed in the passion fruit protoplasts; 2) the optimal method was found to be inoculation of 3 μg plasmid DNA for 5 min with PEG concentration of 40% at the protoplast density of 100 × 10^4^ cells/ml (Fig. 2, 3); 3) Using this method, a high transformation efficiency of 83% was achieved in passion fruit cotyledon protoplasts (Fig. 3B).

**Figure 4.**
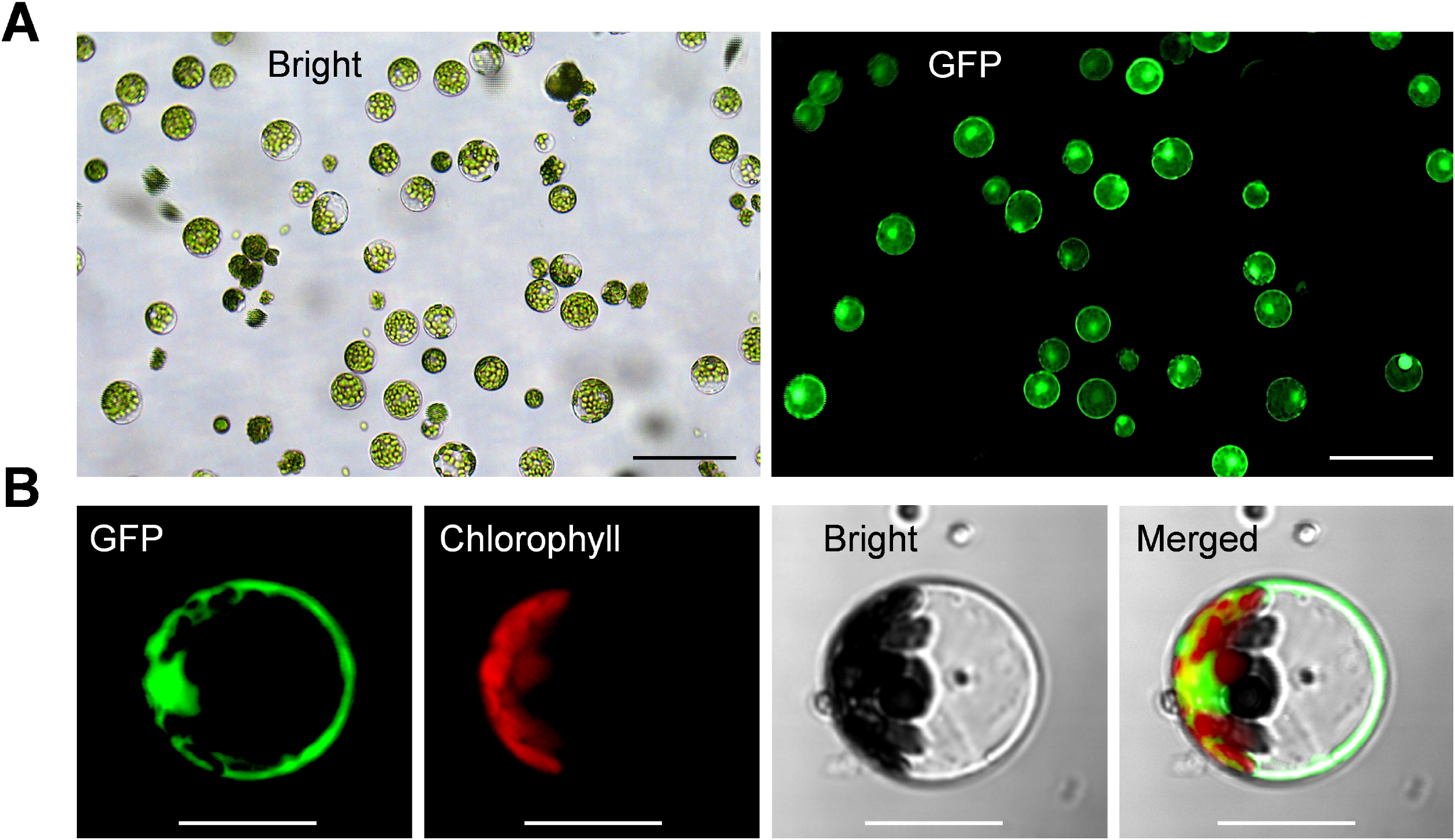
Highly efficient PEG-mediated transformation of passion fruit protoplasts. **(A)** Highly efficient PEG-mediated transformation of passion fruit protoplast with a transformation efficiency of 71.7%, transformed with a green fluorescent protein (GFP)-expressing plasmid (pGreen0029-GFP, 6.0 kb) at 24 h post-transfection (hpt). Scale bar, 100 μm; (**B**) Closed view of GFP expression in passion fruit protoplast under a confocal microscope at 24 hpt. Scale bar, 20 μm.

### 3.3 Subcellular localization studies in passion fruit protoplasts

To assess the suitability of passion fruit cotyledon protoplasts for intracellular localization studies, we used two fluorescent organelle markers for protoplast transformation, including a nucleus marker (H2B-RFP) and an endoplasmic reticulum marker (ER-mCherry-HDEL). At 48 hpt, confocal laser scanning microscopy (CLSM) measurements revealed the correct observation of predicted fluorescence in the corresponding organelles (Fig. 5), suggesting that the cotyledon protoplast serves as an excellent platform for the subcellular localization studies in passion fruit.

**Figure 5.**
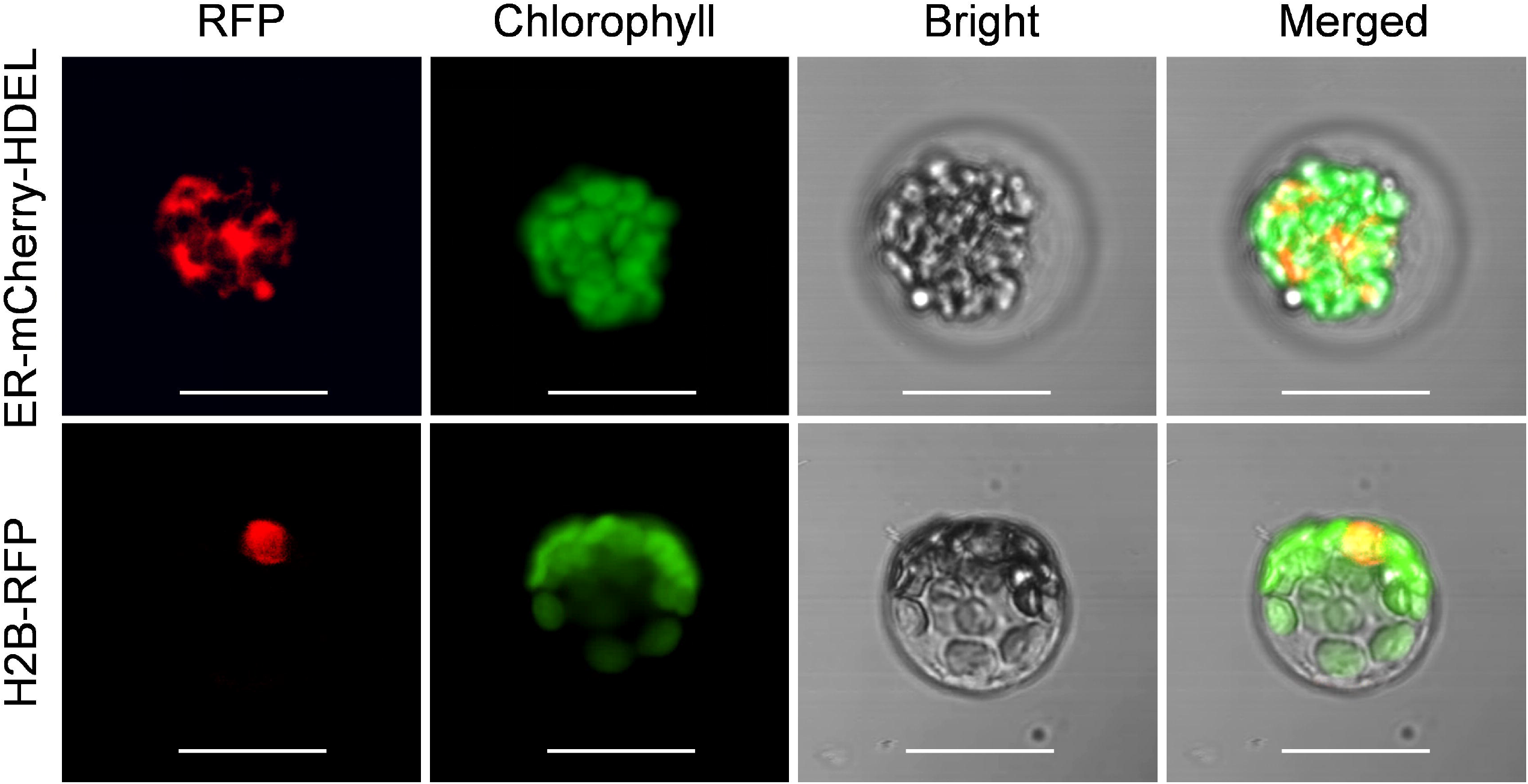
Subcellular localization of various organelle markers. Upper panel: confocal microscopy images of endoplasmic reticulum (ER) maker tagged with mCherry-HDEL at 48 hpt. Scale bar, 20 μm; Lower panel: confocal microscopy images of nucleus maker (H2B protein) tagged with red fluorescent protein (RFP) at 48 hpt. Scale bar, 20 μm.

### 3.4 Protein–protein interaction analysis using the protoplast system in passion fruit

Further, the passion fruit protoplast system was applied to investigate protein-protein interactions using the bimolecular fluorescence complementation (BiFC) assay. The viral coat proteins (CPs) have been well known as capable of self-interactions. In this study, the *cp* gene of telosma mosaic virus (TelMV) was cloned into Gateway-based BiFC vectors (YN and YC), respectively (Fig. 6A) and subsequently delivered into passion fruit protoplasts. As shown in Fig. 6B, co-expression of YN-CP and YC-CP resulted in strong yellow fluorescence throughout the cytoplasm under a confocal microscope, consistent with the CP localization of other potyviruses in intact plant cells (Dai et al., 2020). By contrast, we did not observe any yellow fluorescence in the passion fruit protoplasts transformed with the negative controls YN-CP/YC or YN/-YC-CP. These results demonstrated that the cotyledon protoplast was suitable for protein-protein interaction studies in passion fruit.

**Figure 6.**
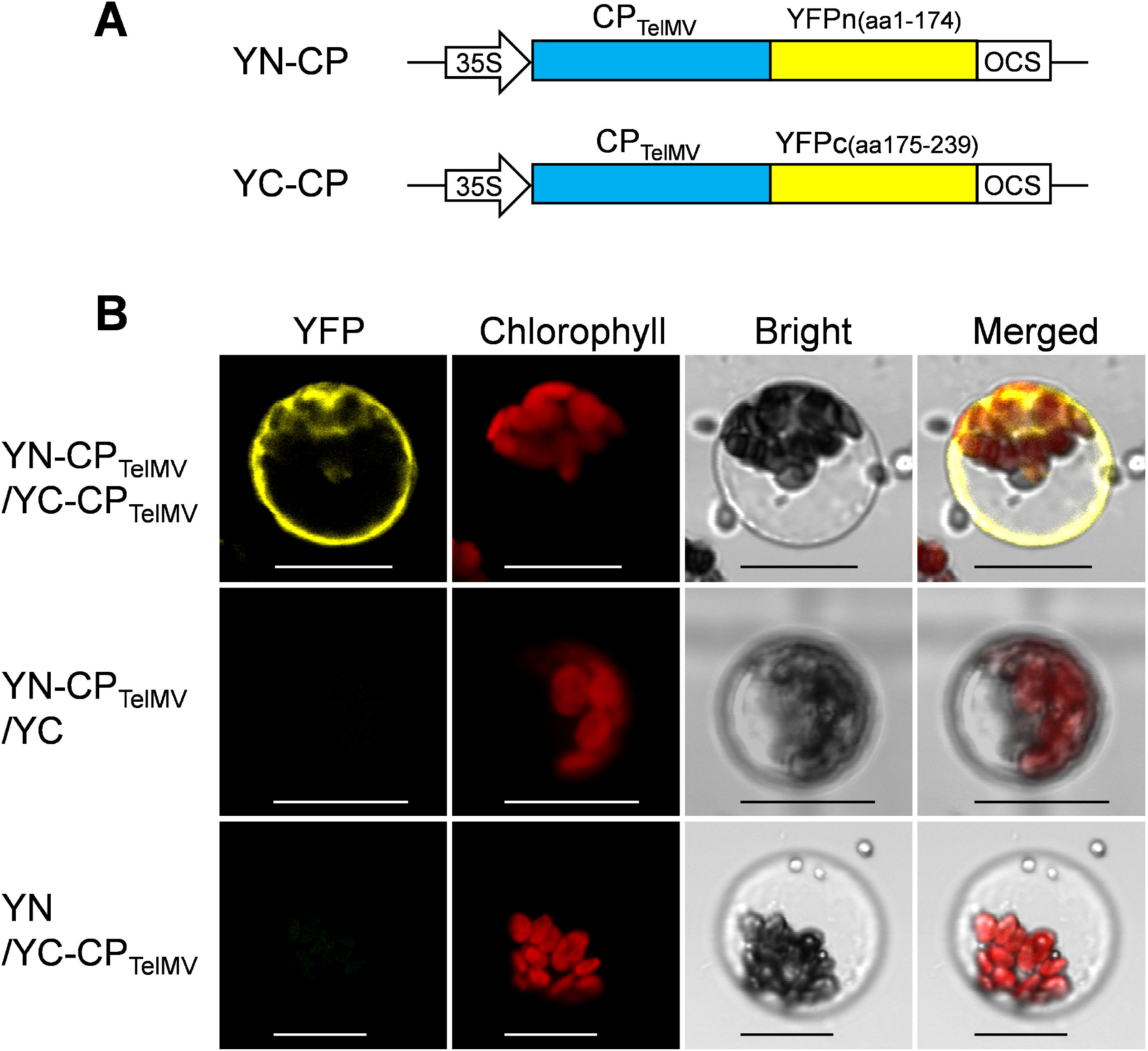
Bimolecular fluorescence complementation (BiFC) analysis of coat protein (CP) of telosma mosaic virus in passion fruit protoplast. **(A)** The schematic illustration of BiFC constructs for viral coat protein expression in passion fruit protoplast; **(B)** BiFC analysis of CP-CP interactions in passion fruit protoplasts. Construct pair of YN-CP/YC-CP was transiently co-expressed in passion fruit protoplasts. Combinations of YN-CP/YC and YC-CP/YN serve as the negative controls. Experiments were repeated three times with similar results. Scale bar, 20 μm.

## 4 Discussion

Herein, we successfully isolated the cotyledon protoplasts from healthy passion fruit seedlings with high yield (2.3 × 10^7^ protoplasts per gram of fresh tissues) and viability (76%) (Fig. 1). With optimization of various transformation parameters, including plasmid amount, protoplast density, PEG concentration and inoculation time, we were able to achieve high transformation efficiency of 83%, delivering a GFP-expressing plasmid DNA into the protoplast through PEG-mediated transformation (Fig. 2-4). Furthermore, the passion fruit protoplast system was successfully applied in subcellular localization and protein-protein interaction studies (Fig. 5, 6).

Traditionally, the leaf serves as the most common source tissue for protoplast isolation of a large number of species, including the model plants Arabidopsis, tobacco, maize, and rice. However, cotyledon has also been applied in successful protoplast isolation for several species, such as cabbage, lettuce, *Malus Pumila Mill* and *Ricinus communis* L (Xu et al., 2021; Ren et al., 2021; Reed and Bargmann, 2021). In the present paper, cotyledon was demonstrated to serve as an excellent source tissue for passion fruit isolation. The protoplast yield was 2.3 × 10^7^ g^−1^FW and is relatively higher than that of *Malus Pumila Mill* (3.72 × 10^6^ g^−1^FW) and *Ricinus communis* L (6.1 × 10^6^ g^−1^FW). The protoplast viability is also comparable to that of the later two species (76% VS 80% and 85%, respectively) (Xu et al., 2021).

In the classic method of protoplast isolation from the model plants Arabidopsis and tobacco, selected leaves were sliced into 0.5-1mm-strips followed by vacuumizing processes (Yoo et al., 2007; Wu et al., 2009). This is also the case for passion fruit, where two independent groups sliced the source tissues into strips for enzyme digestion and protoplast isolation in 1993 (d’Utra Vaz et al., 1993; Dornelas and Vieira, 1993). By contrast, our so-called ‘Cotyledon peeling method’ eliminates the slicing and vacuumizing processes by attaching the cotyledon to the 3M Masking tape with the upper side facing down, and directly peeling away the lower epidermis (Fig. 1). It requires less equipment, time and experimental skills. Furthermore, upon the lower epidermis directly peeling away, it allows the enzyme solution to get much easier access to the mesophyll cells and the intercellular spaces. This method offers great potential for the protoplast isolation of other species in the Passifloraceae family

Previously, plasmid amount has been revealed to affect protoplast transformation efficiency in various species, including grapevine and camellia. The trend of passion fruit transformation efficiency upon adding different amounts of plasmid is raising first and then declining (Fig. 2A, B), which is consistent with that of grape and camellia protoplast. However, the plasmid DNA required for optimal passion fruit protoplast transfection was about twofold lower (Wang et al., 2015; Zhao et al., 2016; Li et al., 2022). In this study, we found that a passion fruit protoplast density of 1.0 × 10^6^ cells/ml resulted in the best transformation efficiency (Fig. 2C, D), which is similar to maize and areca palm (Gentzel et al., 2020; Wang et al., 2023). Different from this, the protoplast density is adjusted to 2-5 × 10^5^ cells/ml in the classic protoplast system of Arabidopsis and tobacco (Yoo et al., 2007; Wu et al., 2009). PEG concentration has been reported to play a major role in protoplast transfection. In our experiment, we found that 40% PEG concentration was optimal for the transformation of passion fruit protoplast (Fig. 3A, B). We have to mention that 40% PEG is the most often PEG concentration used for model plants, including Arabidopsis and tobacco (Yoo et al., 2007; Wu et al., 2009). Our result is also consistent with the detailed optimization of protoplast transformation for multiple species, including *Camellia oleifera* (Li et al., 2022) and Chinese cabbage (Sivanandhan et al., 2021). Different from this, in *Magnolia* and *Freesia* protoplast transformation, 20% was the optimal PEG concentration (Shen et al., 2017; Shan et al., 2019). The difference could be due to different species used for protoplast isolation and/or the physiological status of the plants used. The optimal incubation time for protoplast transfection typically ranges from 2 to 40 mins, depending on the plant species. For example, the optimal incubation time for grapevine, *Magnolia* and cassava protoplast was 2 min, 5 min and 10 min respectively. (Zhao et al., 2016; Shen et al., 2017; Wu et al., 2017). In our system, the most effective transfection time for passion fruit protoplast is 5 min (Fig. 3C, D), which is consistent with the optimization results for *Magnolia* (Shen et al., 2017), but different from *Camellia oleifera*, Chinese cabbage and cassava (Wu et al., 2017; Sivanandhan et al., 2021; Li et al., 2022).

Tobacco leaves and onion epidermal cells are the commonly used plant materials for the subcellular localization studies of heterogenous proteins (Zhao et al., 2016; Priyadarshani et al., 2018). Alternatively, the protoplasts system provides insights into subcellular localization analysis of homogenous proteins at the single-cell level. In the current work, we demonstrated that the passion fruit protoplast could be exploited for protein subcellular localization studies by showing the correct subcellular localization of organelle markers, including the nucleus and endoplasmic reticulum in passion fruit protoplasts (Fig. 5). Future experiments include the investigation of the roles of various passion fruit proteins by subcellular localization and colocalization studies using the established protoplast-based transient gene expression system in passion fruit.

Protein-Protein studies using protoplasts have been broadly used in Arabidopsis and tobacco. Here, we also show our protoplast transient gene expression system could be used for protein-protein studies in passion fruit, illustrated by the BiFC experiments to study two pairs of fusion proteins: YN-CP and YC-CP. A clear YFP signal was observed throughout the cytoplasm upon co-expressing YN-CP and YC-CP (Fig. 6). This result is consistent with its interaction status in the epidermal cells using tobacco leaf (data not shown). Our system provides a vital tool for protein-protein studies *in vivo* when dealing with passion fruit proteins.

## 5 Conclusion

In summary, our study presents the first report of a simple and efficient passion fruit protoplast isolation protocol by the direct cotyledon-peeling method, and establishment of the highly efficient PEG-mediated transformation system upon optimizations. Furthermore, we demonstrate its detailed usage in transient gene expression studies for the first time in passion fruit, including subcellular localization and protein-protein interaction studies. The established protoplast system would provide an essential platform for various passion fruit biology studies, including whole plant regeneration, transgenic studies, gene function analysis and genome editing.

## Data Availability Statement

The original contributions presented in the study are included in the article/Supplementary Material, further inquiries can be directed to the corresponding author/s.

## Author Contributions

ZD, HC, and LW designed the experiment and wrote the manuscript. LW and ZD performed the experiments. All authors analyzed, discussed the data, read, and approved the final manuscript.

## Acknowledgments

We thank Dr. Ji Li (Nanjing Agriculture University, China) for providing pGreen0029-GFP plasmid.

